# DNA glycosylase NEIL2 prevents *Fusobacterium*-mediated inflammation and DNA damage in colonic epithelial cells

**DOI:** 10.1101/2020.06.11.147454

**Authors:** Ibrahim M Sayed, Anirban Chakraborty, Amer Ali, Aditi Sharma, Ayse Z. Sahan, Debashis Sahoo, Pradipta Ghosh, Tapas K Hazra, Soumita Das

**Affiliations:** Department of Pathology, University of California San Diego; Department of Internal Medicine, University of Texas Medical Branch Galveston; Department of Cellular and Molecular Medicine, University of California San Diego; Department of Pediatrics, University of California San Diego; Department of Computer Science and Engineering, Jacob’s School of Engineering; Department of Medicine; John and Rebecca Moore Cancer Center, University of California San Diego

**Keywords:** *Fusobacterium nucleatum*, enteroid, DNA damage, Base-excision repair, colorectal cancer, Microbe-associated inflammation and cancer

## Abstract

Colorectal cancer (CRC) is the third most prevalent and deadly cancer. Approximately, 15-20 % of CRCs display microsatellite instability (MSI); however, the majority (*80–85*%) of cases are sporadic and known as microsatellite stable (MSS). Several recent studies indicated that infection and uncontrolled inflammation initiate DNA damage and lead to cancer progression. One of the major microbes, *Fusobacterium nucleatum* (*Fn*) is highly associated with CRC, but the role of DNA repair in microbe-associated CRC has been largely unknown. Here we show that NEIL2, an oxidized base-specific DNA glycosylase, is significantly downregulated among all the key DNA repair proteins involved in various DNA repair pathways, after infection of *Fn* with stem-cell-based enteroid-derived monolayers (EDMs) of murine and human healthy subjects. Furthermore, following *Fn* infection, NEIL2-null mouse-derived EDMs showed significantly higher level of DNA damage, including double strand breaks, and inflammatory cytokines.. Murine CRC model also showed downregulation of the NEIL2 transcript and accumulation of DNA damage. Importantly, analysis of publicly available transcriptomic data showed that the downregulation of NEIL2 is specific for MSS compared to MSI CRCs. We thus conclude that the pathogenic bacterial infection-induced downregulation of NEIL2, and consequent accumulation of DNA damage, play critical roles in the progression of CRC.

## 1. Introduction

Colorectal cancer (CRC) is the third most prevalent cancer and the fourth most frequent cause of cancer death in the USA and around the world, according to GLOBOCAN 2018 data and the American Cancer Society, 2017 (*1*). CRC is a multifactorial disease that is affected by genetic, epigenetic and environmental factors, such as dietary lifestyle, physical activity, obesity, smoking, alcohol consumption and alteration of gut microbiota (*2–7*). Epigenetic alterations in CRC include aberrant DNA methylation, chromatin modifications, and noncoding microRNA expression (*8*). For the genetic subtype of CRC, the presence or absence of mutations in the DNA mismatch repair system is well studied. However, this hypermutable subtype or deficient mismatch repair (dMMR) system is responsible for only 15% of CRCs, which is known as microsatellite instable (MSI) CRC (*9*). The defect in the MMR pathway leads to microsatellite instability (MSI) and it is a common etiologic factor for the development of CRC such as hereditary non-polyposis colon cancer (HNPCC), also known as Lynch syndrome (*10*). The remaining 85% of the CRCs do not exhibit mutations in the mismatch repair system and known as proficient mismatch repair (pMMR) or microsatellite stable (MSS) CRCs (*11, 12*).

Bacterial/microbial infection can increase the risk of cancer, and the well-known example is *Helicobacter pylori*-infection induced gastric cancer. The alteration of gut microbes can exert a profound influence on host physiology by inducing chronic inflammatory state, affecting stem cell precursors, and the production of toxic metabolites leading to the development of CRC (*13, 14*). *Fusobacterium nucleatum (Fn)* is another pathogenic anaerobic gram-negative gut microbe that is also associated with CRC cases (*15, 16*). *Fn* resides in the oral cavity and plays a role in the development of oral gingivitis and other periodontal diseases (*17*). Oral feeding of APC^Min/+^ mice with *Fn* increased the colon tumorgenesis in these mice (*16*).

Moreover, high levels of *Fn* in CRC patients’ tissues was associated with poor patient outcome and recurrence of infection in post-chemotherapy (*18*). The pathogenesis of *Fn* infection in CRC is initiated by the interaction of FadA of *Fn* with epithelial cells E-cadherin and activates β-catenin signaling that increases the expression of Wnt and inflammatory genes resulting in colorectal tumor progression (*19*); *Fn targets* TLR4 and MYD88 immune signaling to downregulate miRNA involved in the autophagy pathway to alter colorectal cancer chemotherapeutic response (*18*). A recent study showed that *Fn* caused elevated proinflammatory cytokine production and generation of intracellular reactive oxygen species (ROS), leading to oxidative DNA damage.

Host cells then respond to DNA damage by initiating DNA damage response (DDR) after infection. Failure of DDR can lead to the accumulation of mutation, inducing genome instability, and oncogene activation resulting in cancer (*20, 21*). ROS-induced DNA base lesions are predominantly repaired via highly conserved base excision repair (BER) pathway, which is initiated by DNA glycosylases. (*22*). Among five oxidized base-specific DNA glycosylases, OGG1 and NTH1 remove oxidized purines and pyrimidines, respectively, from duplex DNA. *E. coli* Nei-like (NEIL1-3) proteins are a distinct family of DNA glycosylases that remove both purines and pyrimidines. They initiate the excision of lesions present in a single-stranded region, a replication fork, or transcription bubble mimic (*23–28*). Reduced expression of NEIL1, NEIL2, and elevated expression of NEIL3 is involved in the progression of several types of cancer via their association with the somatic mutation load (*29*). NEIL2’s decreased level or loss of its activity, to a less extent, NEIL1, is a risk factor for the progression of cancers such as lung cancer and squamous cell carcinoma in the oral cavity (*30–33*), and poor survival in patients with estrogen-receptor-positive breast cancer (*34*). Recently, we have shown that *Helicobacter pylori-*infection suppresses DNA base-excision repair enzyme NEIL2 and accumulates DNA damage (*35*).

Here we determined the effect of *Fn* infection on the host DNA repair pathways, analyzed the expression level of key proteins using stem cell-based enteroid-derived monolayer (EDMs). Among the DNA repair proteins, MSH2/MSH3 (involved in MMR pathway) and Ku 70 (involved in NHEJ) were upregulated, and only NEIL2 (BER) is downregulated following infection. *Fn* infection increased the inflammatory cytokines, and induced DNA damage in the colonic EDMs. Consequently, *Fn* infection leads to a double-strand DNA break, which was more prominent when BER protein (NEIL2) was depleted. This study demonstrated that *Fn* infection exerted a differential effect on different DNA repair pathways in the colonic EDMs and indicated the critical role of NEIL2 in DNA damage accumulation that linked to CRC following infection. Our results thus provide a novel insight on the link of NEIL2 following *Fn*-infection in downregulating inflammation, DNA damage, and impaired double-strand break repair that can initiate CRC progression.

## 2. Materials & Methods

All experiments involving human and animal subjects were performed following the relevant guidelines and regulations of the University of California, San Diego, and the National Institutes of Health (NIH).

### 2.1 Animals

The *Neil2* knock out (K.O.) C57BL/6 mice were generated and characterized as previously described (*23*). The breeding and maintenance of these mice were carried out as per the approved guidelines of the Animal Care and Ethics Committee of UTMB, Galveston, TX (Hazra protocol 0606029D). APC^min^ mice were bred, maintained, and subsequently used for multiple experiments following the University of California San Diego Institutional Animal Care and Use Committee (IACUC) policies (Protocol #S18086). *All methods were carried out following relevant guidelines and regulations, and the experimental protocols were approved by institutional policies and reviewed by the licensing committee.*

### 2.2 Bacterial cultures

*Fusobacterium nucleatum (Fn)* (ATCC-25586) was obtained from ATCC. *Fn* was cultured anaerobically at 37°C for 48 hours in the chopped cooked meat media (Anaerobic Systems, Morgan Hill, CA), inside an anaerobic chamber containing anaerobic gas kit generating system (MGC, AnaeroPACK System, Japan). Colon enteroid derived monolayer (EDM) was infected with *(Fn)* at a multiplicity of infection (moi) of 100. *Escherichia coli K12 strain DH10B*, (ATCC-PTA¬5105), was cultured on L.B. agar and L.B. broth and used to infect EDM at moi of 100. Adherent-invasive *Escherichia coli* strain LF82 (AIEC-LF82), isolated from the specimens of Crohn’s disease patient, was obtained from Arlette Darfeuille-Michaud and was cultured as previously described (*36*). EDM was infected with LF82 at moi of 100. *H. pylori* strain (ATCC-26695) was incubated with the Brucella Broth supplemented with 10% FBS in a shaker at 37°C in 10% CO2. Colon EDMs were infected with the *H. pylori* at moi 100 for 24 h and 48 h.

### 2.3. Isolation, and maintenance of colonic organoids

Colon organoids were isolated from the colonic tissue specimens of WT C57BL/6, Neil2 KO mice, *CPC-APC*^Min^ mice, healthy human subjects and colorectal cancer (CRC) patients as described previously (*37–41*). Briefly, colon tissues were digested by Collagenase type I [2 mg/ml; Life Technologies Corporation, NY) containing gentamicin (50 μg/ml, Life Technologies Corporation, NY). The digested tissues were incubated at 37°C for 30-40 min cycles with intervals of 10 min each. The tissue specimens were subjected to vigorous pipetting between successive incubations and were constantly monitored to confirm the dislodgement of the intestinal crypts from the tissues. After the separation of 80% of the colon crypts, the collagenase was inactivated with media [DMEM/F12 with HEPES, 10% fetal bovine serum (FBS) (Sigma-Aldrich)], and the colon crypts were filtered using a 70 μm cell strainer. The number of live cells was measured in a hemocytometer following staining by trypan blue (company?). The cells were suspended in the basement membrane matrix (matrigel) (Corning Inc., Kennebunk) supplemented with 50% stem-cell enriched conditioned medium (CM) with WNT 3a, R-spondin and Noggin. Y27632 (ROCK inhibitor, 10 μM) and SB431542 (an inhibitor for TGF-β type I receptor, 10 μM) were added to the media to support the growth of organoids. For the human enteroids, media and supplements were obtained from the HUMANOID CoRE (U.C. San Diego, CA) as done previously (*41*).

### 2.4. The preparation of enteroid-derived monolayers (EDMs)

EDMs were prepared from colonic enteroids as previously described (*36, 42*). Briefly, the enteroids were trypsinized, and the isolated cells were filtered, counted, resuspended in 5% CM, and 2×10^5^ cells (per well) plated in 0.4 μm polyester membrane transwell (Corning, Cat #3470) precoated with matrigel (1:40 dilution)., The cells were then allowed to differentiate for 48-72 h. Transepithelial electrical resistance (TEER) and Lgr5 were used as markers for the quality and differentiation of EDMs (*36, 38*). Differentiated EDMs were challenged with different microbes, as described earlier. The supernatant was collected from the basolateral part of the transwell for cytokine analysis, and the cells were lysed for RNA extraction to check the expression of target genes by qPCR.

### 2.5. RNA isolation and qPCR for gene expression analysis

RNA was extracted from uninfected EDMs, *Fn*-infected and EDMs infected with Fn, E. coli K12-infected EDMs, and LF82-infected EDMs using Quick-RNA MicroPrep Kit (Zymo Research, USA) according to the manufacture’s instruction. cDNA was synthesized using the qScript™ cDNA SuperMix (Quantabio). Quantitative PCR (qPCR) was carried out using 2x SYBR Green qPCR Master Mix (Biotool™, USA) for target genes that and the data was normalized to the housekeeping genes 18s rRNA using the ΔΔCt method. Primers were designed using NCBI Primer Blast software and, the Roche Universal Probe Library Assay Design software (Table 1).

**Table 1.**
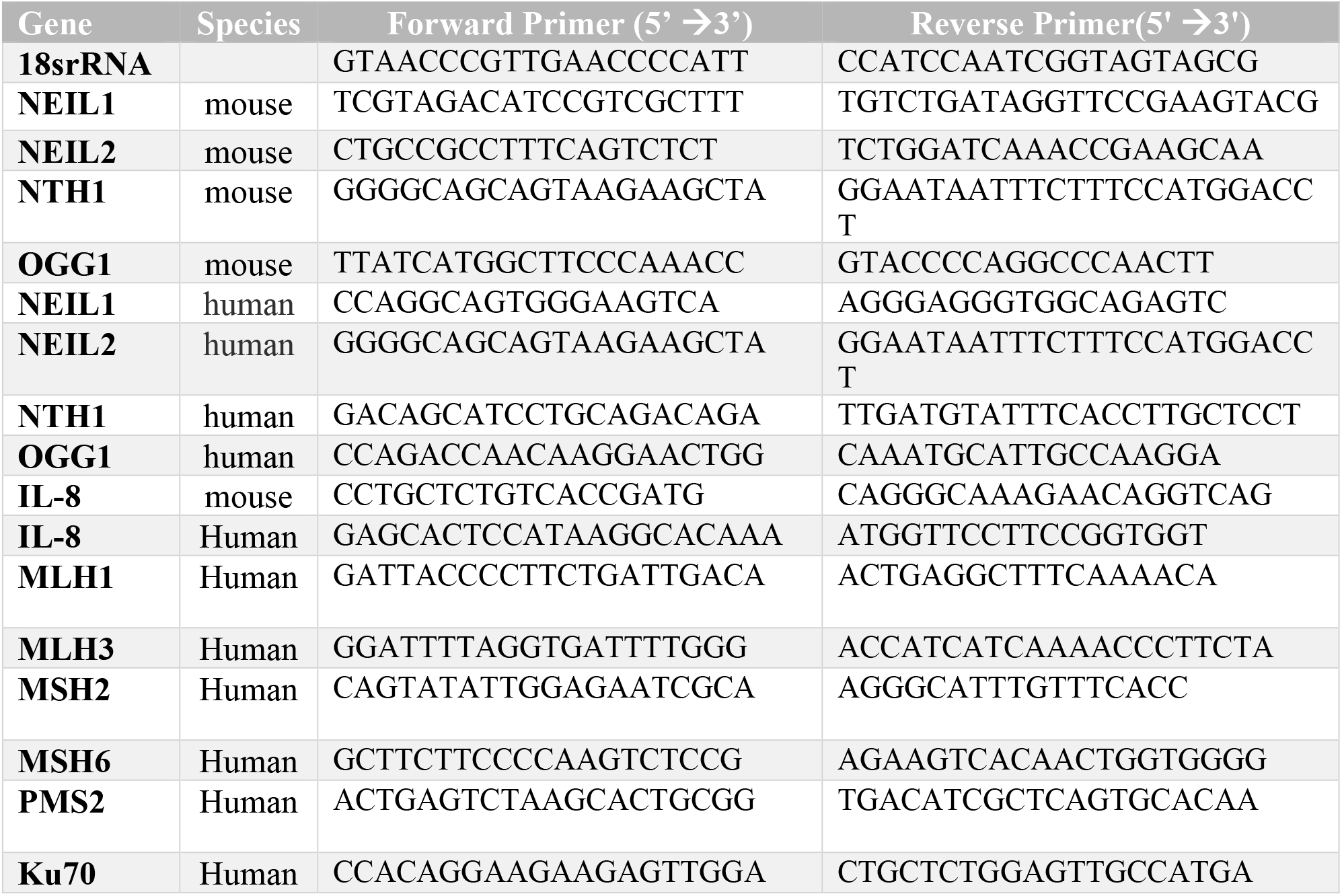

### 2.6. Proteome profiler array

To assess the cytokine levels after *Fn*-infection, a proteome profiler array (Human cytokine array; R&D Systems) was used according to the manufacturer’s instructions. Briefly, the capture antibody of cytokines and chemokines were spotted on nitrocellulose membranes pre-coated with 40 cytokines. The samples /secondary antibody mixture were added to the immobilized capture antibodies on the membrane following the manufacturer’s protocols. After washing, the streptavidin-horseradish peroxidase and chemiluminescent reagents had been added to the membrane. The detected signal was proportional to the amount of cytokine bound. For the quantification of specific cytokines, ELISA was performed.

### 2.7. The measurement of IL-8 cytokine by ELISA

Supernatants were collected from WT., NEIL2 KO EDMs and APC^Min^ polyp either uninfected or infected with *Fn*. IL-8 (K.C.) was measured using the mouse CXCL1/KC DuoSet ELISA kit (R&D Systems) according to the manufacturer’s instructions. The level of IL-8 was compared between *Fn* infected EDMs versus the uninfected EDMs from WT., NEIL2 KO mice and APC^Min/+^ polyp.

### 2.8. Immunoblot for detection of base excision repair proteins and cellular signaling proteins

Whole Cell lysates were prepared with the RIPA buffer (50 mM Tris [pH 7.4], 150 mM NaCl, 1% NP-40, 0.25% Na-deoxycholate, 1mM EDTA with protease inhibitor cocktail [Sigma]), isolated from uninfected, WT-*Fn* infected EDMs, and NEIL2 KO *Fn* infected EDMs, were tested for expression levels of BER proteins using respective antibodies (NEIL2 (in-house) (*23*), NTH1 (in-house) and OGG1 (in house), all antibodies used in 1:500 dilution). GAPDH (GeneTex) was used as loading control. The proteins in the whole-cell extracts from EDMs were separated onto a Bio-Rad 4-20% gradient Bis-Tris Gel, then electro-transferred on nitrocellulose (0.45 μm pore size; G.E. Healthcare) membrane using 1X Bio-Rad transfer buffer. The membranes were blocked with 5% w/v skimmed milk in TBST buffer (1X Tris-buffered Saline, 0.1% Tween 20), then immunoblotted with appropriate antibodies as described above. The membranes were extensively washed with 1% TBST, followed by incubation with anti-isotype secondary antibody (G.E. Healthcare) conjugated with horseradish peroxidase in 5% skimmed milk at room temperature. Subsequently, the membranes were further washed three times (10 min each) in 1% TBST, developed using ECL™ Western Blotting Detection Reagents (RPN2209, G.E. Healthcare) and imaged immediately. For the detection of pATM and total ATM, cell lysates were separated on 8% SDS-PAGE and transferred to PVDF membranes (Millipore). Membranes were blocked with PBS supplemented with 5% nonfat milk or bovine serum albumin and then incubated sequentially with primary and secondary antibodies. Infrared imaging with two-color detection and quantification of Western blots was performed according to the manufacturer’s protocols using an Odyssey imaging system (Li-Cor Biosciences). The dilution of the primary antibodies was as follows: anti-p-ATM, 1:1000; anti-ATM, 1-1000, and anti-tubulin, 1-1000 (used as a loading control).

### 2.9. Assessment of ɤ-H2AX phosphorylation by immunofluorescence (IF)

Either uninfected or *Fn* infected EDMs from WT. and *Neil2* KO mice were fixed with 4% paraformaldehyde (PFA) at room temperature for 15 min, washed once with 1X phosphate-buffered saline, permeabilized and blocked with IF buffer (0.1% Triton TX-100, 2mg/ml BSA, in PBS) for 1 h. Samples were then incubated with anti-p-γH2AX (Santa Cruz biotechnology, INC, 1:500). Incubation with the primary antibody was performed at 4C overnight. After three 10 min washes with PBS, the EDMs were incubated with secondary antibodies at a dilution of 1:500 for 60 min at R.T. in the dark. Washes were performed as before, and the membranes were dried and mounted in Gel/Mount anti-fade aqueous mounting medium (Biomedia Corporation) with coverslips. Images were acquired using a Leica CTR4000 Confocal Microscope with a 63X objective. Z-stack images were obtained by imaging approximately 4-μm thick sections of cells in all channels. All individual images were processed using Image J software.

### 2.10. Analysis of DNA damage accumulation by Long Amplicon Quantitative PCR (LAqPCR)

Genomic DNA (gDNA) was extracted from WT., NEIL2KO, and APCMin EDMs, either uninfected or infected with Fn, using Gentra Puregene Cell Kit (Qiagen) following the manufacturer’s instructions. Accumulation of DNA damage/ strand-break was assessed in both transcribed genes (polß and globin) following the protocol, as described previously (25). Briefly, the extracted gDNA was quantified by Pico Green (Molecular Probes). *E. coli* enzyme Fpg (New England Biolabs) was added to induce the strand breaks at the sites of unrepaired oxidized DNA base lesions. The gene-specific DNA damage strand break accumulation was measured by Long Amplicon quantitative-PCR (LA-qPCR) to by amplifying a 6.5-kb region of polß and the 8.7-kb region of globin genes, according to the following conditions: 94°C for 30 s (94°C for 30 s, 55–60°C for 30 s depending on the oligo annealing temperature, 65°C for 10 min) for 25 cycles and 65°C for 10 min. A small DNA fragment for each gene was also amplified to normalize the amplification of large fragment amplicons, as done before (25, 46). The amplified products were visualized on agarose gels and quantified using an ImageJ automated digitizing system (NIH) based on at least two independent replicates. The extent of damage was calculated in terms of the relative band intensity with WT. untreated subject considered as 100. The information on the oligo sequences is listed in Table 2.

**Table 2.**
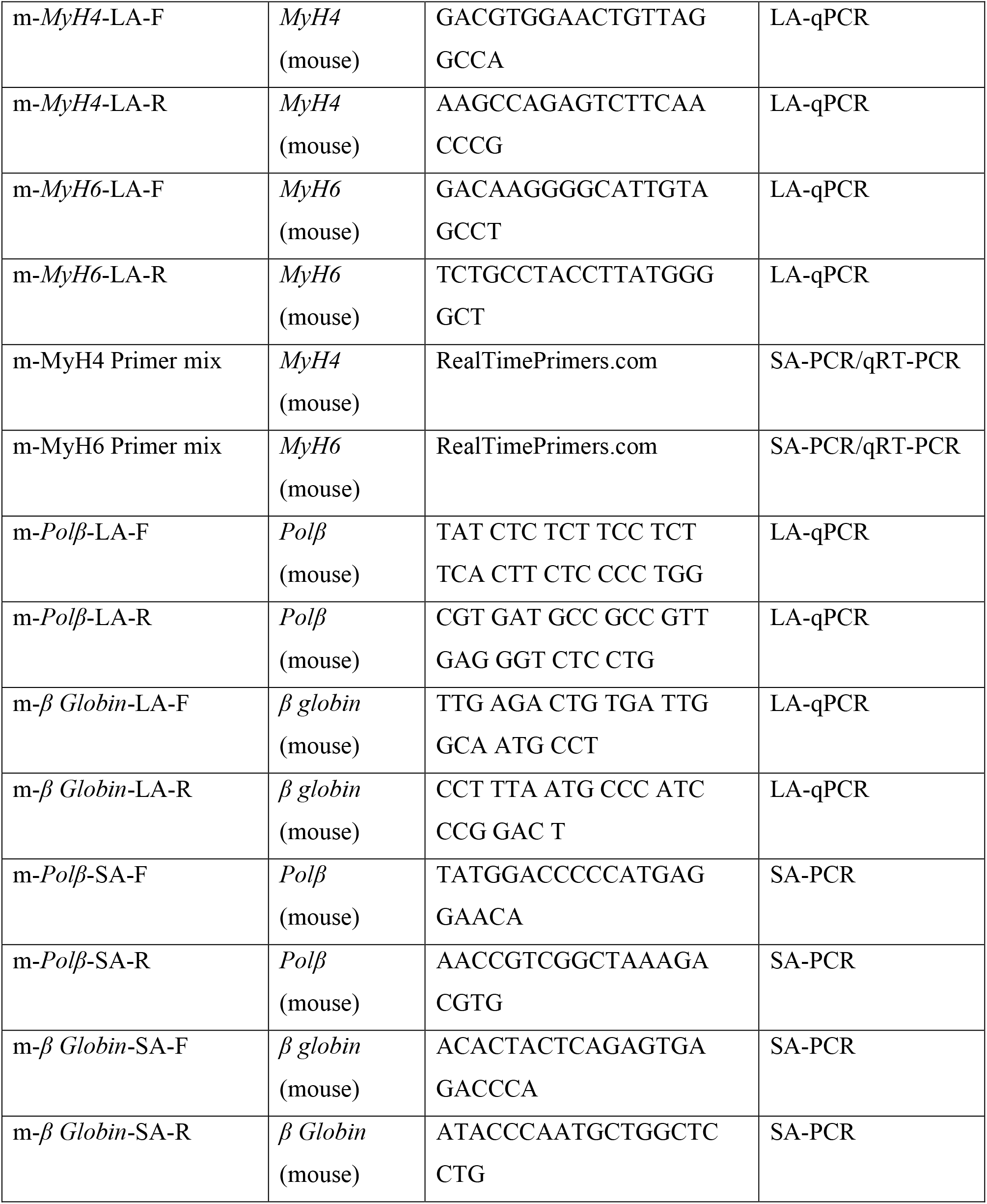

### 2.11. Measurement of oxidative DNA damage

The level of damaged bases was quantified using the DNA/RNA Oxidative Damage ELISA Kit (Cayman Chemical, USA) according to the manufacturer’s instruction and previous papers (*43, 44*). Briefly, this technique is a competitive ELISA assay between oxidatively damaged guanosine bases (8-hydroxy-2′-deoxaguanosine from DNA, and 8-hydroxyguanine from either DNA or RNA) and 8-hydroxy deoxaguanosine acetylcholinesterase conjugate (DNA/RNA oxidative Damage Tracer) for a limited amount of Oxidative Damage Monoclonal antibody (mAb). Since the amount of tracer is constant, the amount of tracer will bind to the mAb will be inversely proportional to the concentration of oxidatively damaged guanine bases in the well. The precoated secondary antibody will react with mAb-damaged base complex and the reaction is developed by the addition of Ellman’s reagent, which is the substrate to acetylcholinesterase. The intensity of the color is measured spectrophotometrically at wavelength 405-420 and it is inversely proportional to the amount of free 8-hydroxy guanosine present in the reactions. According to the manufacturer’s instructions, standard curves were developed using standards provided in the kits and the standards were diluted in the same culture media of infection experiment to confirm that the matrix of standards and the samples is the same. We used this assay to compare the level of oxidatively damaged bases in WT. and NEIL-2 KO colon EDMs either uninfected or *Fn* infected. Also, we used this technique to assess the level of DNA damage generated after infection with several bacteria (*E.coli* K12, *AIEC LF-82*, *H.pylori*, and *E. coli* NC101).

### 2.12. Statistics

Results are expressed as mean ± SEM. *P*-value was determined by GraphPad Prism software 6 (GraphPad Software, La Jolla, USA) using the unpaired student’s t-test. *P* < 0.05 was considered significant.

## 3. Results

### 3.1 NEIL2 is downregulated in the human colonic EDMs following Fn infection

To assess the expression levels of BER enzymes (**Figure 1A**), we infected colonic EDMs with *Fusobacterium nucleatum* (*Fn*) as a model pathogenic bacteria that is known to have a significant association with CRC (*15, 16*). It was found that *Fn* infection significantly downregulated NEIL2 transcript while the NTH1 transcript was upregulated, and the transcript levels of NEIL1 and OGG1 remained unaltered (**Figure 1B**). Since the mismatch repair (MMR) is involved in MSI CRCs, we thus examined the levels of MMR transcripts (*45, 46*). The level of MLH1 and MLH3 remain unchanged after infection, whereas the level of MSH2 and MSH6 were increased, and PMS2 was downregulated following *Fn* infection (**Figure 1C**). Since *Fn* infection caused a decrease in the Ku70 level (a Classical-NHEJ marker) in oral cancer cells (*47*), we tested its level as well in colon EDM and found the upregulation of the Ku70 occurred following *Fn* infection (**Figure 1D**). These results thus demonstrate that *Fn* infection suppressed specifically the DNA glycosylase NEIL2, compared to all other repair proteins tested that are involved in various repair pathways. To demonstrate the specificity of *Fn*-mediated downregulation of NEIL2, human colonic EDMs were infected with commensal *E. coli*-K12 strain (**Figure 1E**) or IBD associated adherent invasive *E.coli* (*E. coli* LF82 isolated from IBD patient) (*36*) (**Figure 1F)**. The transcript level of NEIL2 was not affected following infection with these bacteria (**Figure 1E-1F**), indicating the downregulation of NEIL2 following *Fn* infection is specific.

**Figure 1:**
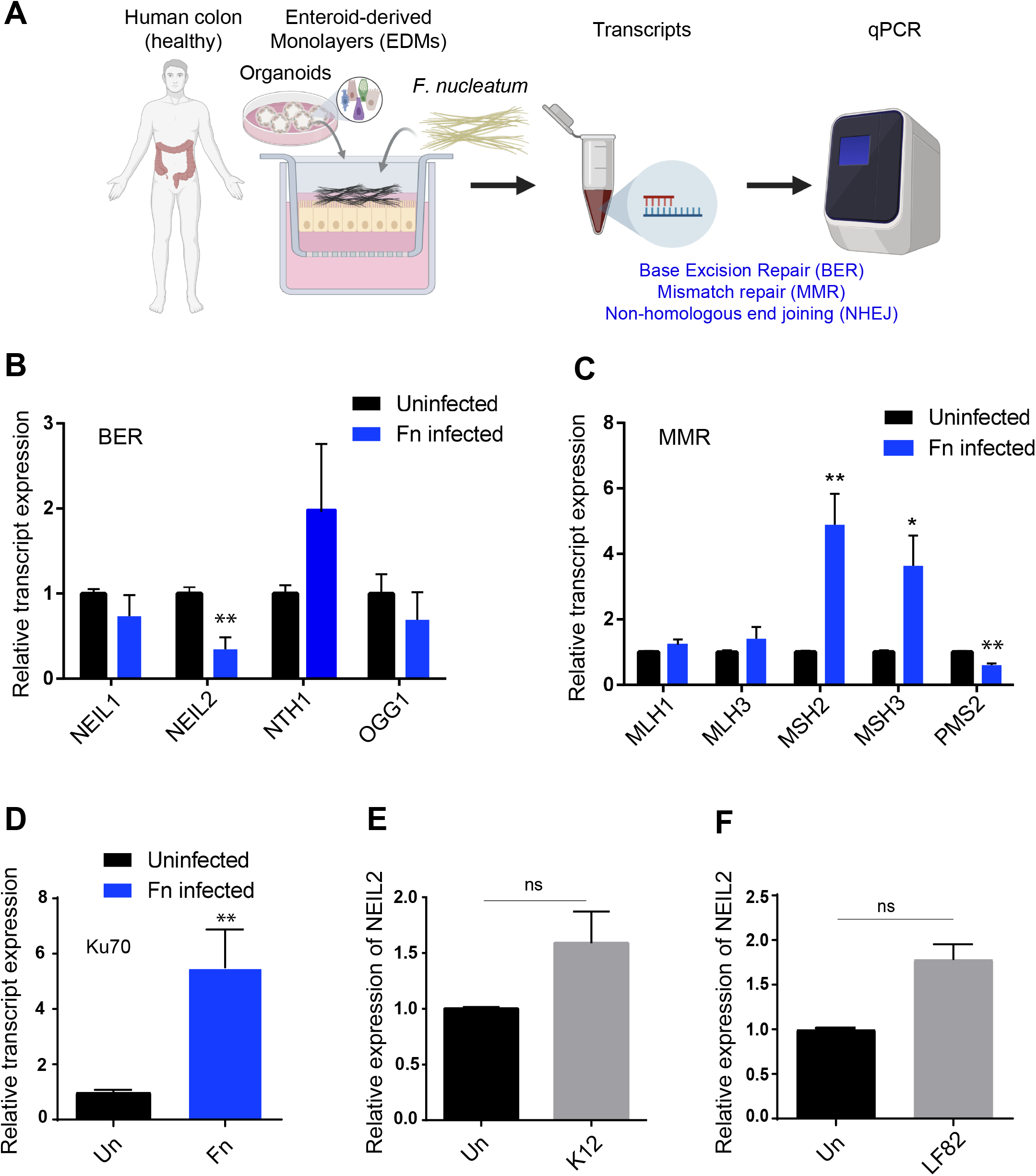
The downregulation of NEIL2 in human colonic EDMs following *Fusobacterium nucleatum* (*Fn*) infection. (A-D) Human colonic EDMs were infected with *Fn* at moi 100 for 24 h. The RNA from EDMs was used for qRT-PCR to determine the expression of genes involved in base excision repair, mismatch repair and for non-homologous end joining (NHEJ). (A) Schematic showing the experimental design. (B) The level of BER transcripts, NEIL1, NEIL2, NTH1, OGG1, (C) The level of MMR transcripts, MLH1, MLH3, MSH2, MSH6, PMS2, (D) The transcript level of NHEJ marker Ku70 were determined by qRT-PCR. (E-F) Human colonic EDMs were infected with commensal *E. coli*-K12 strain (E), or pathogenic IBD-associated adherent invasive *E. coli* LF-82 (F) to determine the expression level of NEIL2 following infection. In (B-F), the expression level of the transcripts was normalized to the housekeeping gene (18srRNA), and the normalized expression value was compared with the respective uninfected control cells. Data represent the mean ± SEM of three separate experiments. * indicates p≤0.05, and ** indicates p≤0.01 as calculated by the unpaired two-tailed student’s t-test.

### 3.2 *Fn* infection suppressed the expression of NEIL2 in the murine colonic EDMs

To determine the effect of *Fn* infection on the expression of BER proteins, we infected murine colonic EDMs with *Fn* and then measured the transcript and protein level of DNA glycosylases by qPCR and W.B., respectively (Figure 2A). Similar to human EDMs, *Fn* infection led to downregulation of NEIL2 and NEIL1 transcripts, while OGG1 and NTH1 transcript levels were either unchanged or upregulated, respectively, probably as a compensatory host mechanism due to *Fn* infection (**Figure 2B**). Next, we assessed the expression level of BER proteins in the EDMs cell lysate., and found that *Fn* infection suppressed the expression of NEIL2 protein. In contrast, the expression of NTH1 and OGG1 were upregulated (**Figure 2C-2D**).

**Figure 2:**
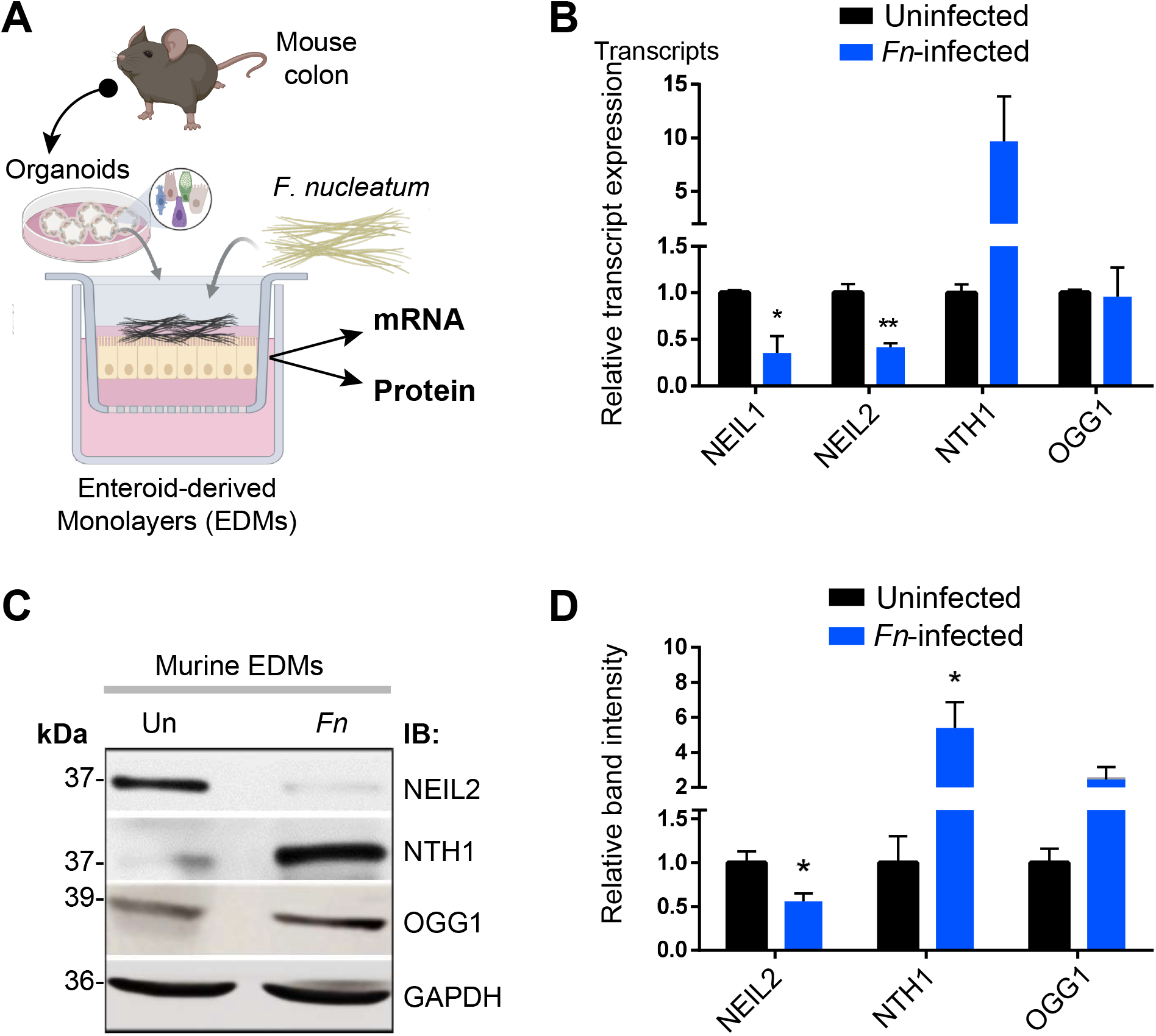
The downregulation of NEIL2 in murine colonic EDMs following *Fusobacterium nucleatum* (*Fn*) infection. **(A) Schematic showing the experimental design. (B)** Wild type (WT.) murine colonic EDMs were infected with *Fn* for 24 h, and the RNA followed by qRT-PCR was performed to detect the level of BER enzymes, NEIL1, NEIL2, NTH1, and OGG1. The expression level of the transcripts was normalized to the 18srRNA and compared with the corresponding uninfected level. Data represent the mean ± SEM of three separate experiments. * and ** indicate p< 0.05 and p≤0.01, respectively, as assayed by the student’s t-test. (C) The level of NEIL2, NTH1, and OGG1 proteins were determined in the whole-cell extract by W.B. in uninfected and *Fn*-infected EDMs (upper panel). The relative band intensity was normalized to GAPDH (loading control) and compared between *Fn* infected and uninfected, which was set as unity (n=1) (lower panel). (D) The relative band intersity of (C) was calculated. * indicate p< 0.05 as assayed by the student’s t-test

### 3.3. The inflammatory response and DNA strand-break produced by*Fn* infection in colonic EDMs depend on NEIL2

To determine whether *Fn* infection can induce inflammatory cytokines, we collected supernatants from colonic EDMs, either mock-infected or infected with *Fn*. By analyzing the proteome profile array, we found that IL-8 and its homolog, CXCL1, were the major cytokines induced and secreted by *Fn-*infected EDMs (**Figure 3A**). The effect of NEIL2 on the expression of pro-inflammatory cytokines was further assessed using the WT. vs NEIL2 KO mouse EDMs following infection with *Fn.* It was found that *Fn* infection upregulated K.C. (IL-8) transcript in both the EDMs; however, NEIL2 KO EDMs had a 4 fold higher K.C. compared to the WT. EDMs following *Fn* infection (**Figure 3B**). Similarly, ELISA analyses also revealed that NEIL2 KO EDMs have a significantly higher K.C. expression compared to the WT. EDMs after *Fn* infection (**Figure 3C**). As we have shown previously that NEIL2 prevents DNA damage accumulation following infection with *H. pylori* (*35*), we, therefore, examined the level of oxidized DNA bases in WT. vs NEIL2-KO EDMs. *Fn* infection indeed led to the accumulation of oxidatively damaged bases, and *Fn* infected NEIL2 KO EDMs accumulated relatively higher level of oxidized DNA base lesions compared to *Fn*-WT EDMs (**Figure 3D**). Also, the level of DNA strand-break was assessed using LA-qPCR in representative Polβ and β-globin genes in the genomic DNA extracted from colonic EDMs from WT. and NEIL2 KO mice after challenge with *Fn*. We indeed observed a significantly higher level of DNA strand-break accumulation in both the Polβ and β-globin genes, in NEIL2 KO EDMs compared to WT. EDMs following *Fn* infection (**Figure 3E**).

**Figure 3:**
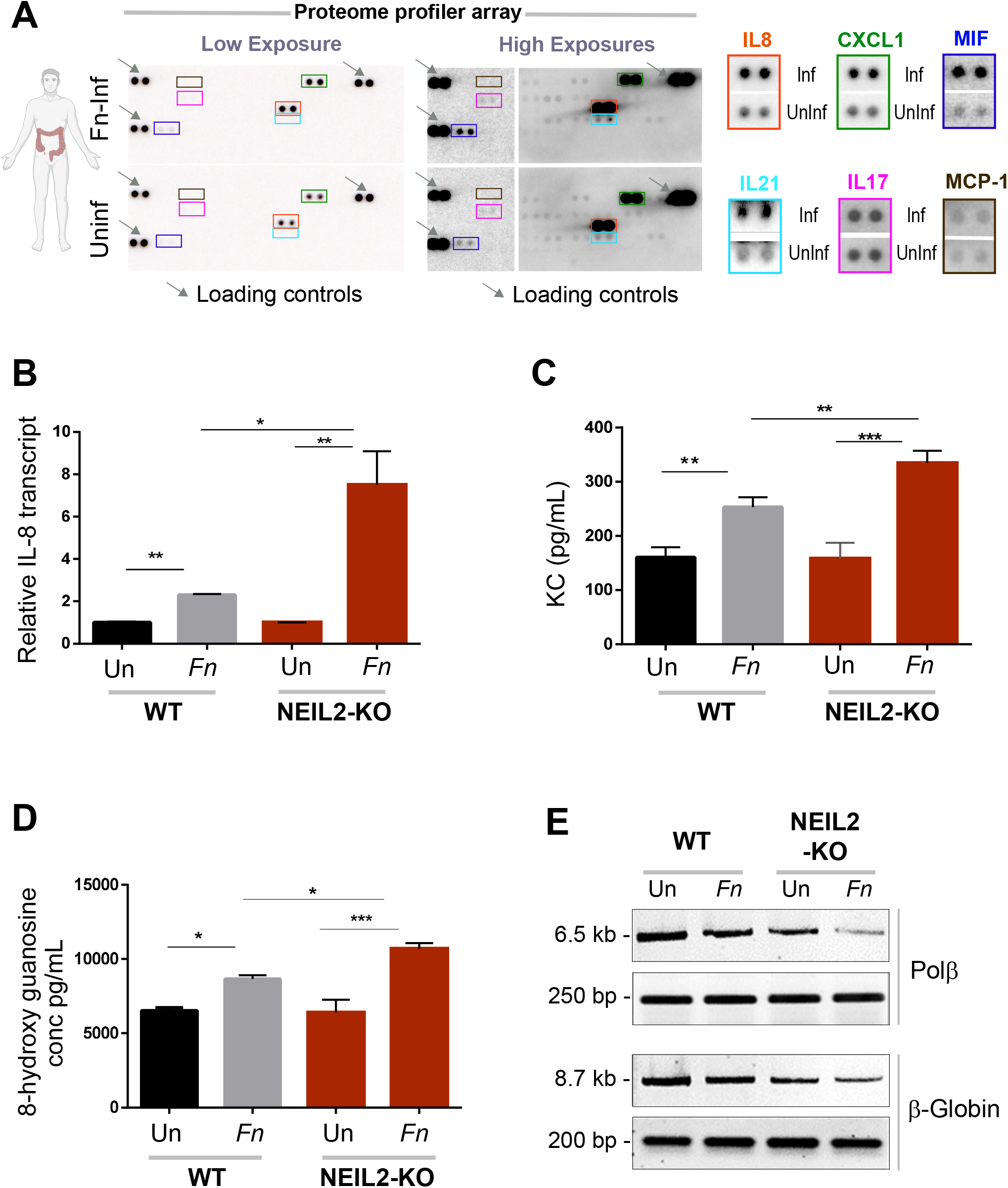
NEIL2 is important for limiting the inflammation and DNA damage generated by *Fn* infection. **(A)** Supernatants from uninfected and *Fn*- infected EDMs were analyzed for cytokines by proteome profiler arrays. Both low (Left) and high (Right) exposures were presented. IL-8 and its homolog, CXCL1 is the major cytokines that were significantly induced in response to *Fn* infection, as shown in the boxes. (B) The differential expression of IL-8/ Cxcl-1 was determined by qRT-PCR of RNA samples isolated from WT. and NEIL2 KO EDMs, either untreated or infected with *Fn*. (C) The level of Cxcl-1/KC cytokine was measured in the supernatants from the EDMs of WT. or NEIL2 KO mice, either uninfected (Un) or infected with *Fn* by ELISA. (D) The supernatants of WT. and NEIL2 KO EDMs, either uninfected (Un) or infected with *Fn*, were analyzed for oxidative DNA/RNA damage. (E) WT. and NEIL2 KO EDMs were either uninfected or infected with *Fn*, then harvested for genomic DNA isolation. LA-qPCR was performed to evaluate the level of oxidized DNA base lesions after *Fn* infection. Representative gels show the amplification of each long fragment (~7-8 kB) (upper panel) normalized to that of a short fragment (~250 bp) (lower panel) of the corresponding (Pol β and β-Globin) genes. Data was generated from the mean ± SEM of three independent experiments where * p<0.05, ** p<0.01 and *** p<0.001 as assayed by the t-test.

### 3.4. NEIL2 knockdown exacerbated*Fn* induced DNA double-strand breaks (DSBs)

As *Fn* infection increased DNA strand-break, we next checked whether it could induce DNA double-strand break. We thus investigated the formation of p-γH2AX foci, a marker of DSBs, by confocal microscopy, and found that *Fn* infection induced the level of p-γH2AX (**Figure 4Ai**). We then assessed the possible link between NEIL2 deficiency and DSB accumulation following *Fn* infection and found significantly higher p-γH2AX expression in NEIL2 KO EDMs compared to WT. EDMs following *Fn* infection (**Figure 4A**). Only 30% of cells in WT. EDMs showed the formation of p-γH2AX foci whereas 70% of cells in NEIL2 KO EDMs showed p-γH2AX foci after *Fn* infection (**Figure 4B**).

**Figure 4:**
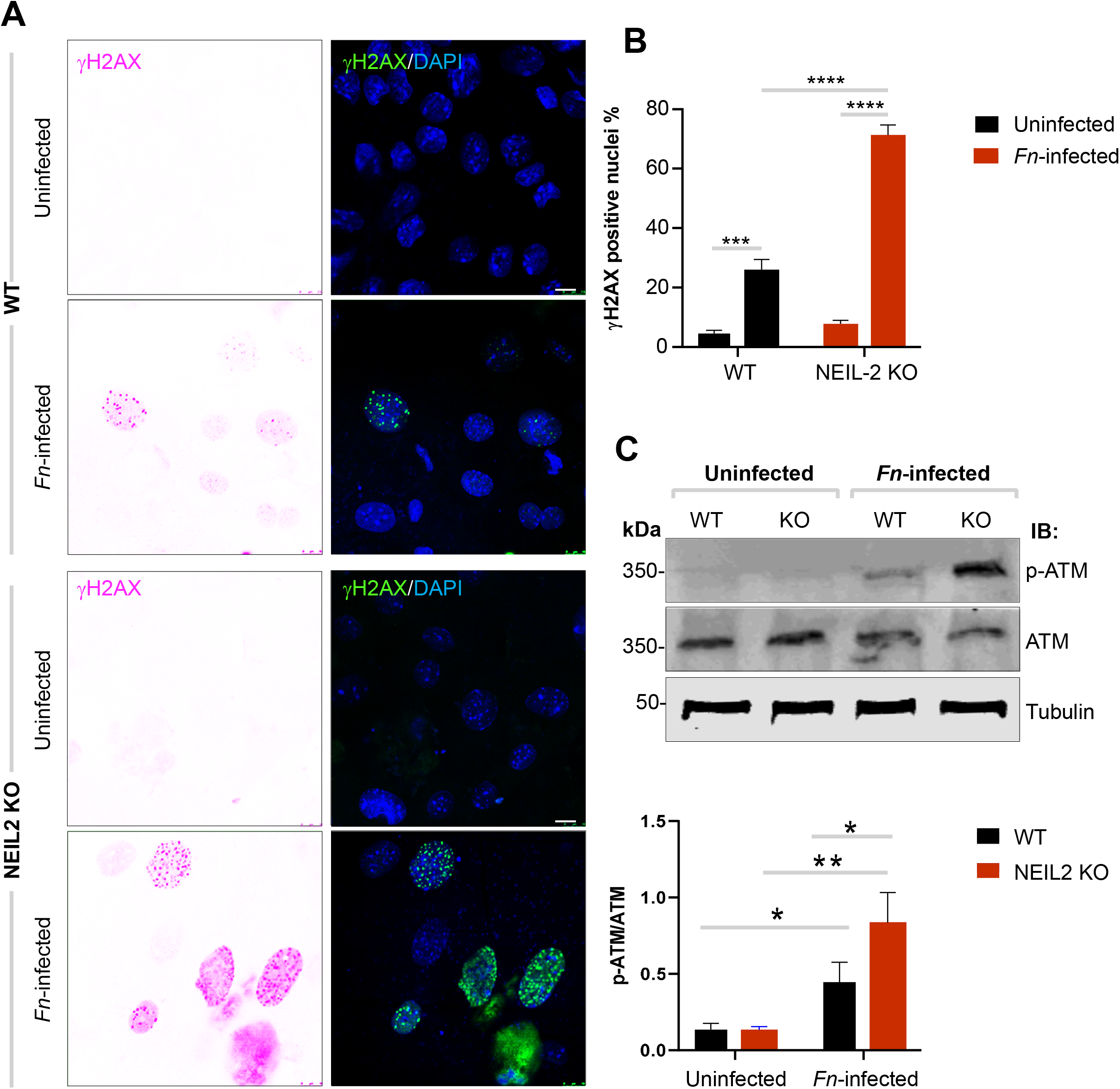
NEIL2 knockdown exacerbated *Fn* induced double-strand DNA breaks (DSBs). (A) The level of p-γ-H2AX (a marker of DSBs, green) was determined by IF, DAPI was used for nuclei staining (blue). WT. and NEIL2 KO colonic EDMs either uninfected or infected with *Fn* were used to measure the amount of DSBs. (B) The percentage (%) of the p-γ-H2AX positive nuclei was calculated. (C) The protein levels of phosphorylated ATM (pATM) and total ATM was determined in both EDMs by W.B. Tubulin was used as a loading control. The relative protein expression levels of pATM normalized with those of total ATM were quantified by the Image J software. The expression level of pATM/ATM was compared in *Fn*-infected EDMs vs uninfected EDMs, and *Fn-*NEIL2 KO EDMs vs *Fn*-WT EDMs. Data was generated from the mean ± SEM of three independent experiments. * indicates p<0.05, ** p<0.01, *** p<0.001, and **** p<0.0001 as determined by the student’s t-test.

Next, we assessed the effect of *Fn* infection on the activation of Ataxia telangiectasia mutated kinase (ATM) protein by Western analysis. ATM is a serine/threonine kinase that regulates cell cycle checkpoints and DNA repair. ATM is activated by autophosphorylation (pATM) on Ser1981 in response to DSBs (52). We indeed found that Fn infection increased the expression of pATM protein, and the ratio of pATM/total ATM was significantly higher in *Fn* infected EDMs than the corresponding uninfected EDMs. Also, the expression of pATM was significantly higher in *Fn* infected NEIL2 KO EDM compared to *Fn* infected WT EDMs (Figure 4C), indicating DNA DSB accumulation did occur in NEIL2-deficient cells.

### 3.5. *Fn* infection downregulated NEIL2 and increased oxidative damage in the murine CRC model

To understand the effect of *Fn* infection on the expression of BER proteins in CRC, we analyzed the most commonly used CDX2 Cre APC^Min/+^ mouse as a murine CRC model. These mice developed multiple intestinal neoplasia and died within 8-12 weeks and are used to simulate human familial adenomatous polyposis (FAP) (53). Also, *Fn* infection can increase the colon tumorgenesis in these mice in vivo (16). The enteroids were isolated from the uninvolved/unaffected region (normal, no polyp) and the involved/affected regions (polyps) of the colonic specimens of CDX2 Cre APC^Min/+^ mice. Similar to our previous observation in humans and WT murine EDMs, *Fn* infection suppressed the transcription of NEIL1 and NEIL2. In contrast, the transcription of NTH1 and OGG1 was either upregulated or unchanged in EDMs generated from uninvolved and involved regions (**Figure 5A-B**). *Fn* increased the inflammatory response in APCMin/+ polyp EDMs, as shown by the increase in KC/IL-8 expression by qRT-PCR and by ELISA (**Figure 5C-5D**).

**Figure 5:**
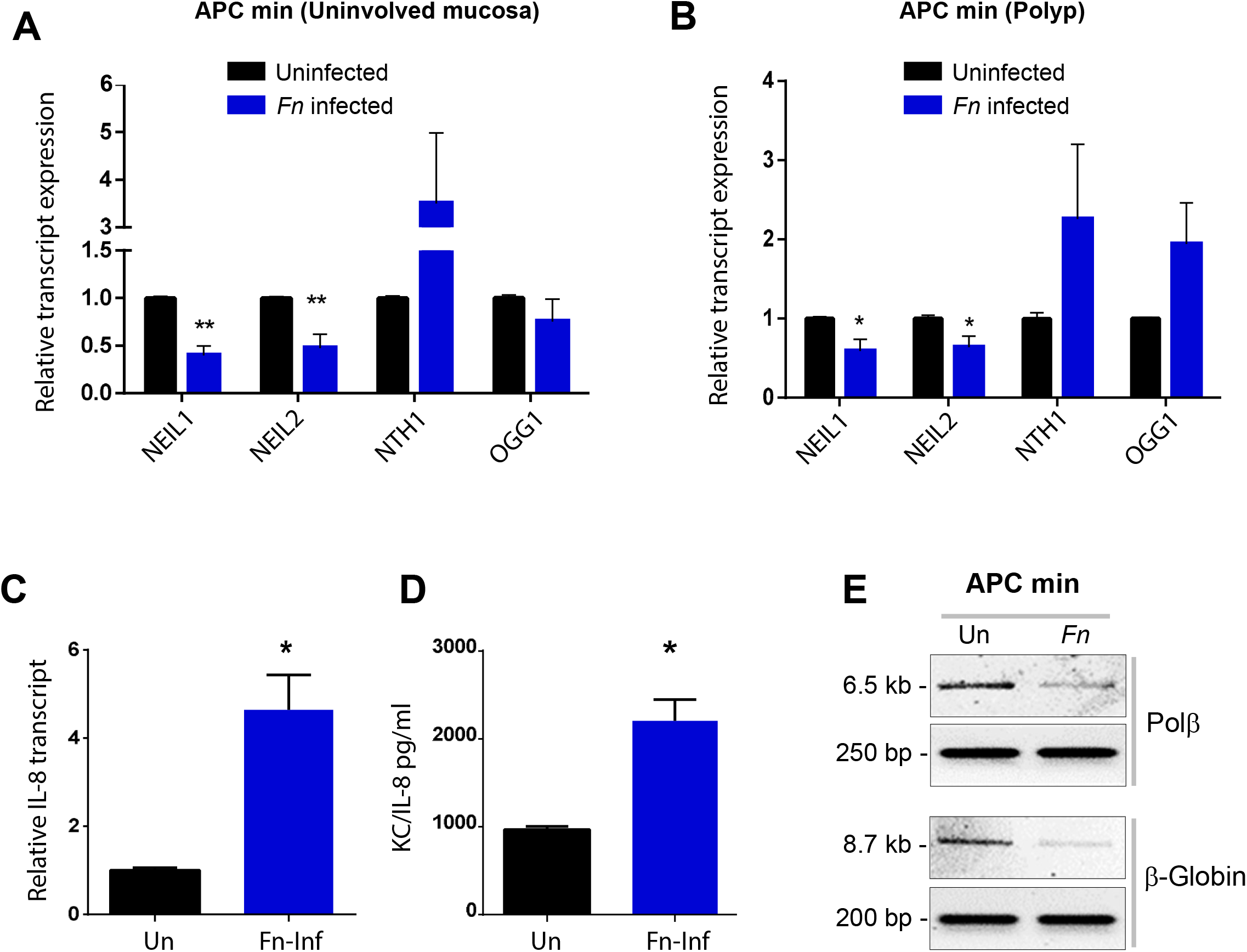
*Fn* infection downregulated the NEIL2 transcript and increased the oxidative damage in the murine CRC model. (A) APC^Min/+^ EDMs derived from the uninvolved region of the colon were infected with *Fn*, then the level of BER transcripts (NEIL1, NEIL2, NTH1, OGG1) was determined by qRT-PCR. Data represent the mean ± SEM of four separate experiments. ** indicates p≤0.001 as assayed by the student’s t-test. (B) APC^Min/+^ EDMs derived from polyp region of the colon were infected with *Fn*, then the level of BER transcripts (NEIL1, NEIL2, NTH1, OGG1) was determined by qRT-PCR. Data represent the mean ± SEM of three separate experiments. * indicates p≤0.05 as assayed by the student’s t-test. (C) APC^Min^/+ EDMs derived from polyp region of the colon were infected with *Fn*, then the transcript and expression level of KC/IL-8 was determined by qPCR (D) The level of Cxcl-1/KC cytokine was measured in the supernatants from the EDMs of APC^Min/+^ EDMs derived from polyp region, either uninfected (Un) or infected with *Fn* by ELISA (E) LA-qPCR was performed to evaluate the level of oxidized DNA base lesions after *Fn* infection in APC^Min/+^ EDMs. The Representative gels show the amplification of each long fragment (~7-8 kB) (upper panel) normalized to that of a short fragment (~250 bp) (lower panel) of the corresponding (Pol β and β-Globin) genes.

Next, we assessed the DNA strand-break levels in the genomic DNA extracted from APC^Min/+^ EDMs, either untreated or infected with *Fn* using LA-qPCR for Polβ and β-globin genes (**Figure 5E**). We observed a significantly higher level of DNA strand-break accumulation in both the Polβ and β-globin genes after *Fn* infection compared to uninfected EDMs (**Figure 5E**). To determine the impact of *Fn* infection on oxidative damage generated in APC^Min/+^ EDMs, we measured the level of oxidatively damaged bases in the supernatants of APC^Min/+^ EDMs infected with *Fn* and compared with commensal *E. coli* K12 and other gut pathogens associated with CRC or IBD (**Figure 6A**). A significantly higher level of oxidative DNA base damages was produced by colon cancer-producing bacteria such as *E.coli* NC101 strain, *H. pylori*, and *Fn* (**Figure 6A**) than commensal and other gut pathogens. Comparing the level of oxidized base adducts produced from different colon cancer pathogens compared to untreated EDMs, we found that *Fn* infection had the highest oxidative DNA damages, followed by *H. pylori* and *E. coli* NC101 (**Figure 6B**).

**Figure 6:**
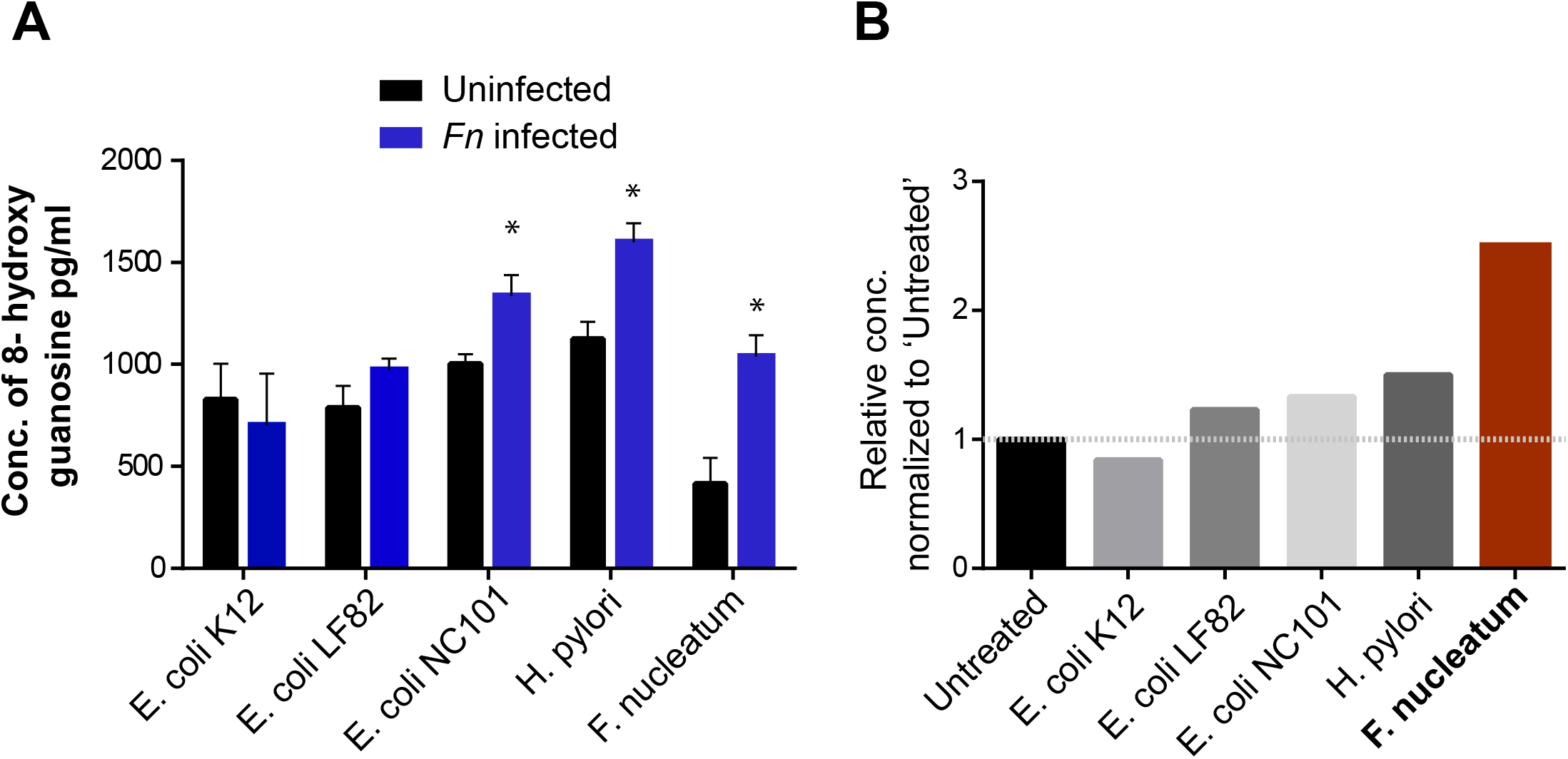
*Fn* infection was the highest inducer of the oxidative DNA/ RNA damage in the murine CRC model. (A) APC^Min^/+ EDMs derived from the uninvolved region of the colon were infected with different microbes; commensal *E. coli* K12, IBD-associated adherent-invasive *E.coli* LF82 and colon cancer-associated pathogens (NC101, *H. pylori* and *Fn*). The supernatants were collected from the uninfected and infected EDMs done in the same experiments and assessed for oxidative DNA damage (right). Data represent the mean ± SEM of three separate experiments. * indicates p≤0.05, ** indicates p≤0.01 as assayed by student’s t-test. (B) The relative level of the oxidized bases produced by each microbe was compared with uninfected cells, which is considered as 1. The relative production of the oxidized base was compared between different microbes

Collectively, these results showed that Fn could suppress the expression of BER proteins NEIL2 in murine CRC model, leading to an increase in the inflammatory response, and accumulation of oxidized DNA base lesions, which could eventually lead to DNA strand-breaks and thereby contribute to the development of CRC.

### NEIL2 is downregulated in the MSS-CRCs patients

DNA damage response (DDR) alterations are found in 17 % cases of G.I. cancers, including CRCs (*48*). CRC is classified into two categories: i) The microsatellite instability-high (MSI-H) group, with defects in the DNA mismatch repair (MMR) system and accounts for 15% of tumors; and ii) the microsatellite stable (MSS) group, which exhibits chromosomal instability and accounts for the remaining 85% of tumors (*49–51*). Previously, a set of gastric cancers with and without microsatellite instability (named MSI and MSS, respectively) showed the differential expression profile of some of the DNA repair genes (*52*). Here, we have examined the expression of NEIL2 in MSI vs MSS CRCs in four different publicly available datasets with GSE13067, GSE13294, GSE26682, and GSE18088 (*53–57*). Interestingly, the analysis of samples from the MSI vs MSS group revealed that the level of NEIL2 is significantly decreased in the MSS group compared to the MSI group (**Figure 7**). We did not notice a significant difference between the two groups in the level of other BER proteins, such as NEIL1 and OGG1 (**Figure 7**). These results thus suggest that NEIL2 plays a significant role in *Fn*-induced MSS-CRCs.

**Figure 7:**
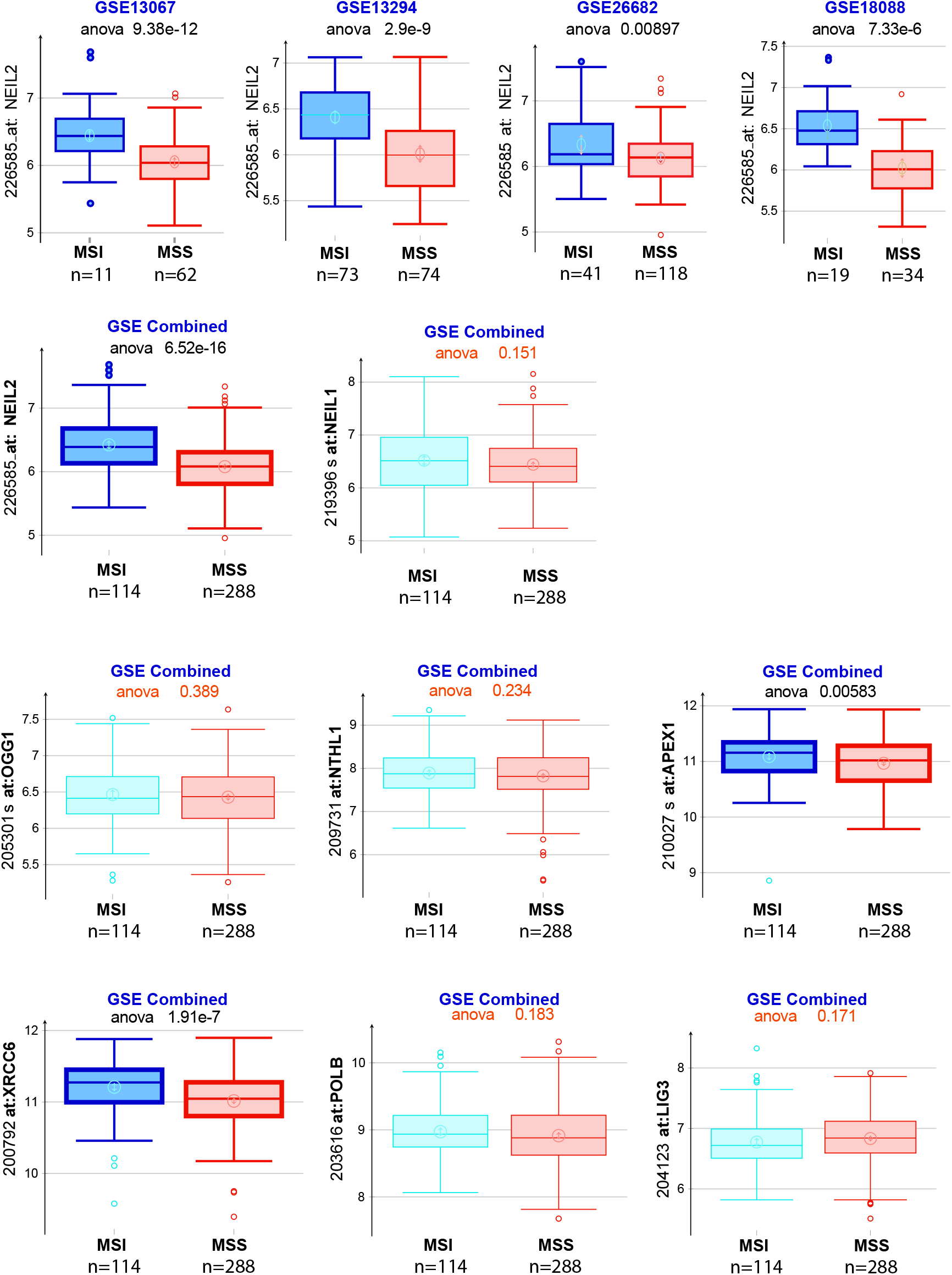
Analysis of NEIL2 in CRCs patients. The expression of BER proteins was analyzed in MSI and MSS groups from the different publicly available datasets GSE13067 (*83*), GSE13294, GSE26682 (*55*), GSE18088 (*54*).

## Discussion

Persistent microbial infections and chronic inflammation represent major risk factors for cancer development. Several pathogens cause chronic inflammation and induce DNA damage via the release of reactive oxygen and nitrogen species (RONS), which eventually leads to genomic instability (*58*). *In vivo* studies showed that BER is the main pathway that repaired the DNA damage induced by chronic inflammation and reduced the risk of colon carcinogenesis in mice (*59*). Deficiency of BER enhances the inflammation-associated colon tumorigenesis and increases the risk of development of gastric cancer following *H.pylori* infection (*59*). Recently, we have shown that NEIL2 is downregulated after *H. pylori* infection in gastric epithelial cells and causes DNA damage (*60*).

Similar to *H. pylori* in gastric cancer, *Fn* is associated with CRC (*61*). *Fn* is detected in 10-90% of CRC patients, with higher prevalence in the proximal colon than distal colon (*61–63*). Previous studies have supported a causal effect of *Fn* in CRC (*64, 65*). Binding of *Fn* has unique adhesin, FadA, to E‐cadherin leads to activation of Wnt/β‐catenin signaling, which causes overexpression of Wnt genes, oncogenes c‐Myc, Cyclin D1, and inflammatory genes (*64*). *Fn* selectively promotes the growth of cancerous cells by inducing Wnt/β‐catenin modulator Annexin A1 (*65*). Also, *Fn* promotes chemoresistance to CRC by modulating autophagy (*18*). But to date, there is no report on the link between *Fn*-mediated CRC, DNA repair pathways, and inflammation that can promote cancer progression. This is the first study to our knowledge, where DNA repair pathway and specifically the role of DNA glycosylase NEIL2 has been studied in colonic epithelial cells following *Fn* infection.

Using 3D stem-cell-based organoid-derived monolayer models from human and murine, we found that *Fn* especially suppressed BER protein NEIL2 at both the transcript and protein level. The downregulation of NEIL2 is specific for *Fn* infection but not after the challenge with commensals or other gut pathogens. *NEIL2*-KO EDMs showed higher inflammatory cytokine (KC/IL-8), the higher level of oxidative DNA/ RNA damage after *Fn* infection. The expression of NEIL2 is also downregulated in a murine model of APC^Min/+^ mice with significant DNA damage and higher inflammatory cytokines. *NEIL2*-KO EDMs showed higher levels of p-γH2AX and p-ATM after *Fn* infection and connected NEIL2 with DSBs. A recent study showed that *Fn* promoted the oral squamous cell carcinoma (OSCC) by upregulating the expression of γH2AX and accumulating DSBs, leading to accelerating the cell cycle and increasing the cell proliferation capacity (*66*). Similarly, *H. pylori-*activated ATM due to the induced host genomic instabilities produced by cag PAI-positive strains through the accumulation of DSBs (*67*). Previous work has shown that reduced expression of NEIL2 protein is associated with the progression of several types of cancer, including CRC (*29*). Also, lower NEIL2 expression within each stage of the disease led to a lower probability of survival compared to patients with higher NEIL2 expression (*52*). Our results showed that *Fn* infection specifically suppressed NEIL2, which in turn increases inflammatory response and DNA damage leading to the initiation and progression of CRC (as summarized in **Figure 8**).

**Figure 8.**
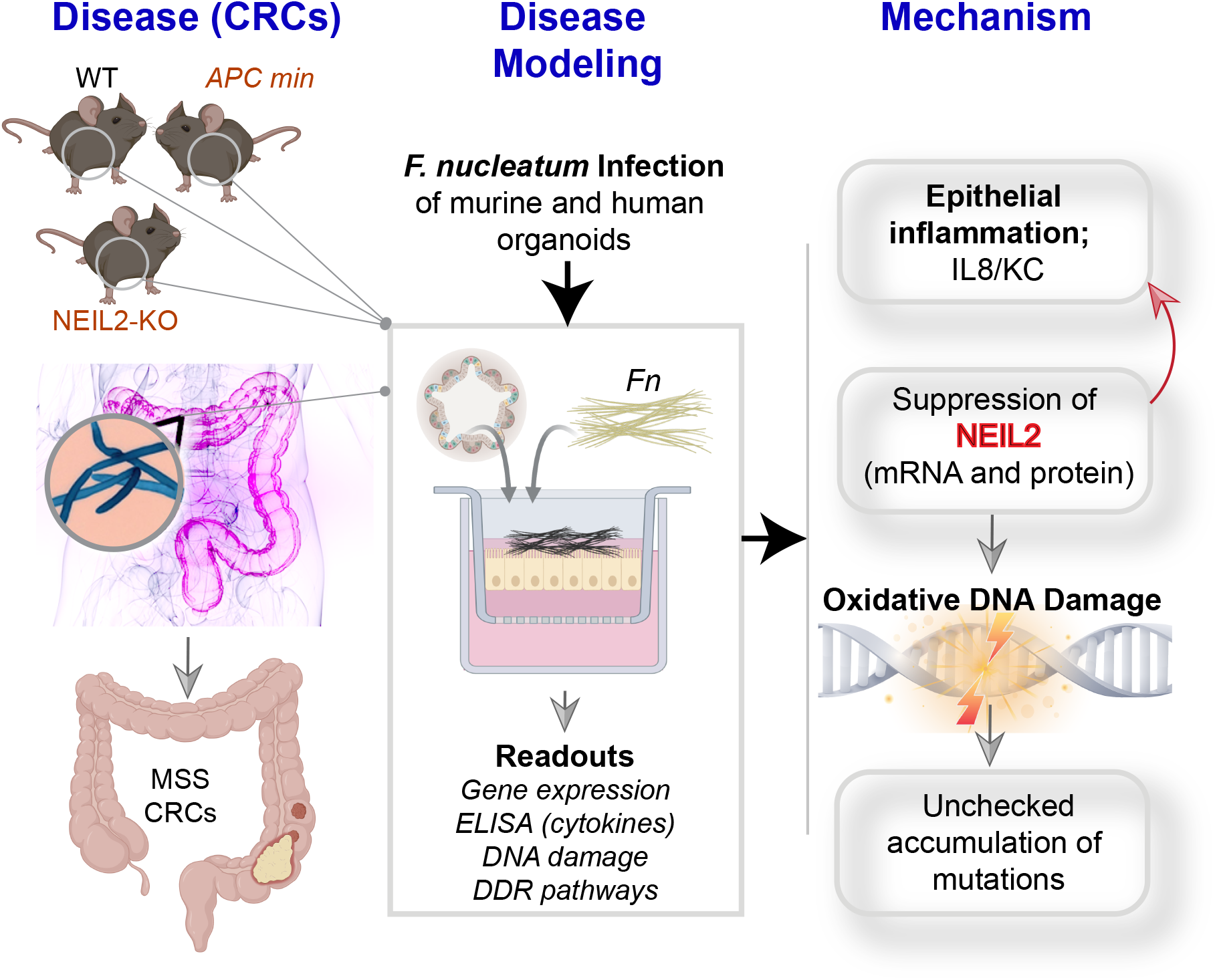
Schematic Model. The model is showing the infection of *Fn* to human and mouse EDMs. *Fn* infection specifically suppressed NEIL2; consequently, the inflammatory response and DNA damage will be exacerbated, leading to the initiation and progression of CRC.

Our study with APC^Min/+^ mice is an excellent experimental model for studying genetic, epigenetic, environmental, and therapeutic aspects of intestinal neoplasia in humans (*68*). Oral feeding of *Fn* to APC^Min/+^ mice resulted in accelerated small intestinal and colonic tumorigenesis, infiltration of specific myeloid-derived immune cell types into tumors, and a similar of NF-κB pro-inflammatory gene signatures as human (*64*). Other bacterial pathogens have been identified that promote colitis-associated CRC in APC ^Min/+^ mice such as *Bacteroides fragilis* and adherent-invasive *Escherichia coli* strain NC101 (*69, 70*). We used both polyp and noninvolved (N.I.) regions of APC ^Min/+^ and *Fn* infection downregulates BER transcripts of NEIL2, inflammatory response, and DNA damage.

One of the major sources of oxidative DNA damage in the colon is the disbalance of bacterial communities. Bacterial infection-induced inflammation leads to the generation of reactive oxygen species (ROS) that can result in oxidative damage, which is more prominent, especially with bacteria causing chronic infection (*24*). Bacterial species that play a role in the pathogenesis of CRC include *Bacteroides fragilis*, *Helicobacter pylori*, *Escherichia coli*, *Enterococcus faecalis*, and *Clostridium septicum* (*71, 72*). Among all these *Fn* is the most prevalent and associated with more than 50% of CRCs, including MSI and MSS cancers.

### Our finding and role of BER in CRC

Finally, we investigated the level of NEIL-2 transcript in CRC patients, and we found that the NEIL-2 transcript is downregulated in CRC derived organoids compared to healthy subject derived organoids. Also, we checked the gene expression of several BER and related genes in MSI and MSS CRC cohorts using the publicly available data. Interestingly, only NEIL2 is downregulated in the MSS CRC cohort compared to the MSI cohort. MSS accounts for 80-85% of colorectal cancer patients, and it is mainly due to chromosomal instability. While MSI, accounts for 15% of CRC, occurs when the MMR genes stop functioning at their highest potential, areas of DNA could start to become unstable due to the errors. It is often in tumors associated with the hereditary syndrome, Lynch syndrome, and some sporadic cancers.

BER is the main DDR that removes the oxidative and alkylating DNA damaged bases without affecting the double helix DNA structure (*73, 74*). Therefore, BER is essential for maintaining genomic stability and preventing the initiation and progression of cancer (*29, 75*). Genetic defects or Inherited variation in BER proteins is a predisposing factor for the development of CRC (*76, 77*). The direct evidence of the role of oxidative DNA damage and its repair is proven by hereditary syndromes (MUTYH-associated polyposis, NTHL1-associated tumor syndrome), where germline mutations cause loss-of-function in glycosylases of base excision repair, thus enabling the accumulation of oxidative DNA damage and leading to the adenoma-colorectal cancer transition. Unrepaired oxidative DNA damage often results in G:C>T:A mutations in tumor suppressor genes and proto-oncogenes and widespread occurrence of chromosomal copy-neutral loss of heterozygosity. However, the situation is more complicated in complex and heterogeneous disease, such as sporadic colorectal cancer. Previously it has been shown that BER-DRC (DNA repair-capacity) represents a potential prognostic biomarker, and it is associated with patients’ survival and prediction of therapy response (*78*). Base excision repair imbalance in colorectal cancer has prognostic value and modulates Oxidative DNA damage and BER depletion has higher PDL1 expression (*79*). The underlying mechanism needs further investigation.

Our finding of higher DSBs in the *NEIL2* KO mice EDMs following infection with *Fn*, opened a new direction of the involvement of NEIL2 with DSBs. It is known that NHEJ, mediated by DNA-dependent protein kinase (DNA-PK) complex and ligase IV/XRCC4, can sense and bind to any DNA double strand break (DSBs) ends, which makes it the most common DSB repair pathway in mammalian cells. DNA-PK consists of DNA binding ends Ku70/K80 heterodimer, initiates the DDR pathway through activating cell cycle detection points, and activating apoptosis programs, they are predominant in G1 of the cell cycle (*66, 80–82*). Our results showed that *Fn* infection specifically suppressed NEIL2, that in turn increases inflammatory response and accumulates DNA damage and DSBs that can initiate the progression of CRC.

## Acknowledgment

This work was supported, in whole or in part, by National Institute of Health Grants: DK107585, C3 Padre Pedal MCC pilot grant (to S.D.); R01 NS073976 (to T.H.); and R01HL145477 (S.S. and T.H.); W81XWH-18-1-0743 (to S.S. and T.H.); R01 AI141630 (to P.G.); R01-GM138385 (to D.S.) and UG3TR002968 (P.G. and S.D.). We are grateful to the HUMANOID CoRE for providing the organoid media and for biobanking and culture a part of the human organoid.

## Abbreviations

CRC: Colorectal cancer
EDMs: Enteroid-derived monolayers
MOI: Multiplicity of Infection
WT.: Wild Type
BER: Base Excision Repair
DGs: DNA glycosylases
LA-qPCR: Long Amplicon quantitative-PCR

